# Habitat suitability modeling growth and cover trends of staghorn coral (*Acropora cervicornis*) outplants in the lower Florida Keys

**DOI:** 10.1101/2025.07.11.664295

**Authors:** Glenna Dyson, Erich Bartels, Easton R. White, Ian R. Combs, Kristen Mello-Rafter, Thomas C. Lippmann, Jennifer A. Dijkstra

## Abstract

The decline of important reef building corals has motivated the development of habitat suitability models used to identify optimal locations for coral restoration. In the Florida Keys habitat suitability models incorporate coarse spatial data sampled over large areas, resulting in recommended outplant sites at distant locations, making it logistically difficult and expensive to access and regularly monitor. Restoration efforts to date show that outplanting success can vary widely within a limited space, necessitating improved predictive abilities of coral outplant success at high spatial resolutions within a restoration site. With the advent of Structure-from-Motion image reconstruction, fine-scale, site specific, digital terrain models can be created to support habitat suitability model development. In this study, generalized linear mixed models used extracted seafloor terrain attributes and environmental variables to identify within site locations of high *Acropora cervicornis* growth and healthy coral cover of long-term outplants. Percent healthy coral cover significantly decreased after two years of outplantation. The submodel of corals exclusively less than two years old was unable to identify environmental conditions associated with higher healthy cover. For all corals, outplant recommendations for higher healthy cover are in deeper waters, away from the coast, in less rough terrain, and closer to the reef edge. Model results for growth support these recommended outplant sites, in addition to concave locations near high slope relief. Finally, our results also indicate that marine heat waves, but especially marine cold waves negatively correspond with coral growth, and high wind events positively correspond with coral growth. These model results provide a basis for further endeavors in modeling endangered organismal success, which are vulnerable to minute differences in local environmental conditions.

## Introduction

In the past half century, corals around the world have been decimated due to mounting anthropogenic pressures including disease, overfishing, river run off, high intensity thermal and wind events (1–4). These threats have resulted in increased efforts to promote resilient reefs, capable of withstanding and recovering from disturbance through the creation of marine protected areas, genetic engineering of corals towards elevated heat resistance, and outplanting corals. Stony reef-building corals have been the primary target of outplantation. One such genus, *Acropora*, has been targeted due to forming much of the essential reef structure throughout the Florida Reef Tract (5) and its outplantation facilitates variation in benthic coral composition and fish species (6). Despite being a good candidate for outplantation, due to asexual propagation and a fast growth rate (7), the genus has shown to be especially vulnerable to environmental stress (8,9) and disease (10,11). To support efforts towards enhancing reef resiliency, habitat suitability models (HSMs) targeting *Acropora* restoration have been developed, including models of wild *Acropora* population occurrence (12), recruitment (13), and global responses to ocean warming and acidification (14).

These HSMs have the potential to identify optimal sites for *Acropora* outplantation, however models for outplant success have been challenging for coral reef managers and outplanters to implement. Current models often use coarsely resolved environmental data (9), many collected through remote sensing efforts, resulting in recommended sites that are financially and logistically difficult to access for outplantation and recurrent monitoring. Predicted suitable habitat can be hundreds of square kilometers, although many restoration activities occur at much finer spatial scales (10s of meters) where coral survival throughout the same site is highly variable (15). Additionally, current models have primarily used wild *Acropora* populations. Coral outplant’s growth and survivorship perform differently than wild and transplanted coral populations (16,17), making these models imprecise in recommendations for outplants. Lastly, when coral outplants are surveyed, long-term monitoring is often not realized nor influential in informing outplanting techniques, and thus sustained recovery of *Acropora* outplants is not clearly established (18,19). Consequently, there is a knowledge gap in HSMs using fine-scale, within site, terrain attributes for long-term coral outplant locations.

This study aims to develop HSMs utilizing fine-scale terrain attributes which may contribute to within site variations in growth (total and healthy) and healthy cover of a commonly used stony coral for outplantation, *Acropora cervicornis*. *A. cervicornis* has been one of the primary targeted *Acropora* species for restoration due to its ecological importance as a reef builder and its physiological characteristics as a long-lived, fast growing species (20–22). HSMs developed thus far have shown improved wild *A. cervicornis* populations in waters with moderate sea surface temperatures and limited temperature ranges, moderate turbidity to mitigate temperature fluctuations and UV radiation (9), and higher water flow to deliver food and oxygen (9,23). From these results, restoration work has called for outplanting deeper, or where there is higher water flow to decrease the negative impact of high intensity thermal events (24). These HSMs show *A. cervicornis* has limited suitable habitats within the Florida Reef Tract, constrained to the fore and back reef of the lower and upper Florida Keys, the Dry Tortugas, and nearshore Broward-Miami reefs (15,23). This study’s use of fine-scale environmental data aims to delineate optimal coral outplant sites within such regions by modeling outplant growth and percent healthy cover within the forereef of the lower Florida Reef Tract. Growth (separated into total and healthy) indicates where at fine-scales there is optimal mass transport (exchange of nutrients and dissolved gasses) and minimal stress (25,26). Total growth indicates where the most growth is occurring, whereas healthy growth indicates where the most growth is occurring despite infection from disease. Percent healthy cover indicates where corals are least exposed to disease, are more resilient to disease, or where corals are best able to recover from infection (27).

In this study, we use fine-scale three-dimensional terrain attributes derived from Structure-from-Motion (SfM) photogrammetry, environmental data, and high intensity thermal and wind events as predictor variables for modeling *A. cervicornis* growth and percent healthy cover. The use of SfM is unique in these HSMs for its capacity to develop three-dimensional terrain models which provide insight in terrain structure and complexity (28–30); the scale of these models is also several orders of magnitude finer (mm) than spatial data often used in HSMs (0.25km to 1 km). This study aims to address the utility of SfM and fine-scale data to predict site-specific optimal coral outplant locations. Model results contribute towards further understanding of *A. cervicornis* restoration and the applicability of fine-scale models for other endangered species.

## Methods

### Outplanting

Percent cover and growth of *A. cervicornis* were determined at eight outplant sites in the lower Florida Keys. Corals were outplanted by Mote Marine Laboratory at Sand Key, Eastern Dry Rock, M32, ICC1, and ADAC1 (Fig 1). All sites reside within the Florida Keys National Marine Sanctuary. Additionally, Eastern Dry Rock and Sand Key are sanctuary preservation areas.

**Fig 1.**
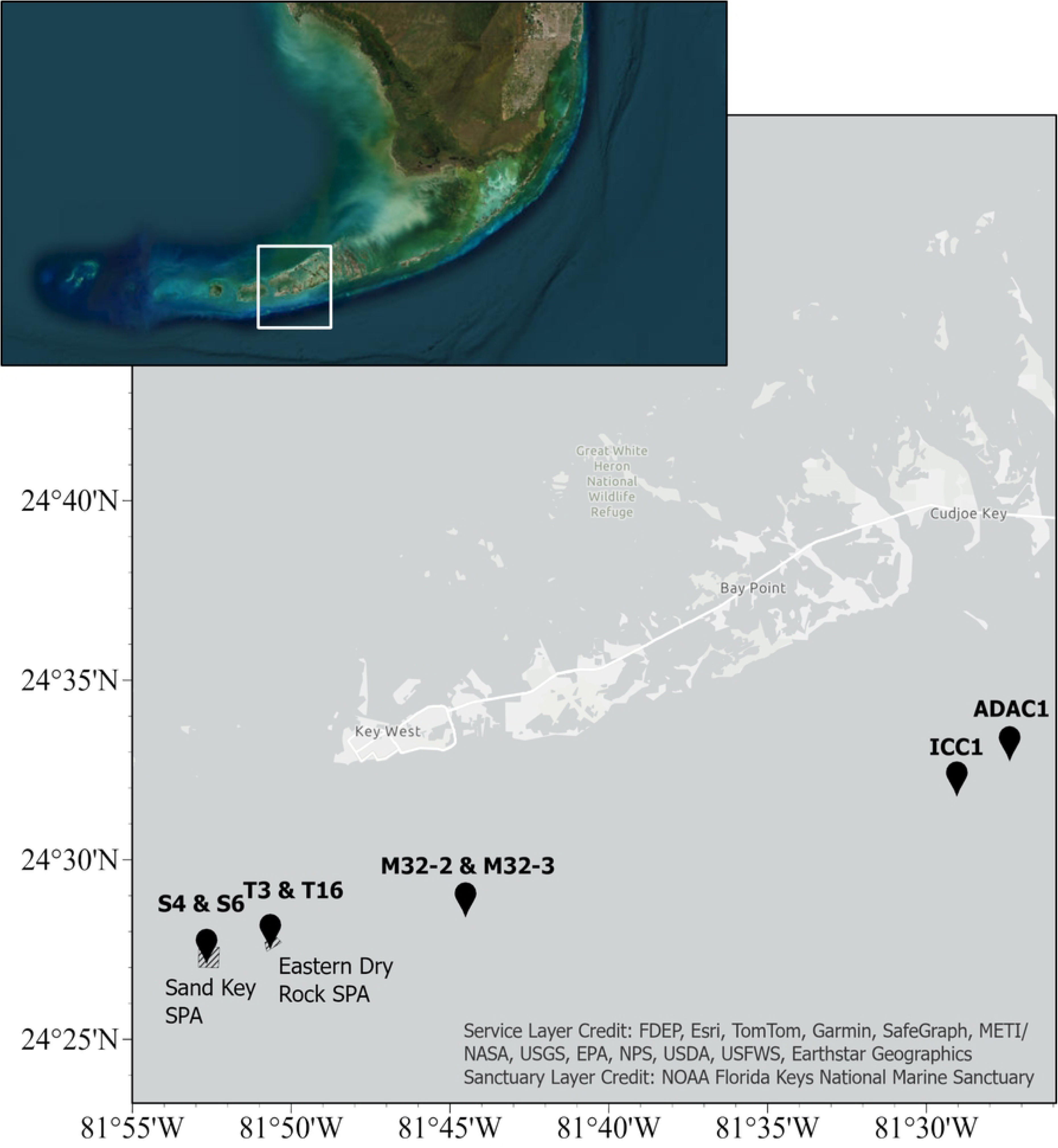
Map of coral outplant sites. Eight survey sites within the lower Florida Reef Tract to west of Key West (S4, S6, T3, T16, M32-2, M32-3, ICC1, and ADAC1). Shaded areas refer to sanctuary preservation areas.

Coral outplant fragments ranged from 250 to 1000 per outplant site, and the corals’ initial sizes ranged from 3cm to 50cm. Except for the singular 3cm coral plugs at ADAC1, all other sites had outplanted clusters of five coral fragments. Corals were affixed to the seafloor using either nails and cable ties; nails, cable ties, and epoxy; or affixed to plugs with epoxy. To enhance genetic diversity, a variety of genotypes were outplanted.

### Image Collection

Eight coral outplant sites were surveyed in June of 2022, and one of the eight sites (ICC1) was previously surveyed the year before in July of 2021, resulting in nine outplant surveys (Table 1). Coral outplant ages ranged from 129 to 1,788 days. All sites, except ADAC1, are on the forereef of the Florida Reef Tract. Images were collected in 10 x 30m plots at all sites. Coded targets (which also functioned as scale bars to ensure accurately sized orthomosaics) were systematically placed throughout each site in a grid formation to support co-registration. Images were taken with two Canon 70D DSLR cameras housed in an Aquatica A70D. Planar images of the seafloor were collected in a lawn-mower survey pattern to facilitate co-registration and limit distortion, resulting in ∼ 80% image overlap. Depth measurements taken from a dive computer accounted for the four corners of each 10 x 30m site.

**Table 1.** Summary of coral outplant locations and time spent outplanted.

| Outplant ID | Location | Outplant Date | Age of outplants when surveyed |
| --- | --- | --- | --- |
| T3 | Eastern Dry Rock | 7/8/2017 | 1788 days |
| S4 | Sand Key | 9/11/2018 | 1386 days |
| S6 | Sand Key | 3/25/2019 | 1191 days |
| M32-2 | Marker 32 | 8/9/2019 | 1050 days |
| M32-3 | Marker 32 | 5/6/2020 | 779 days |
| ICC1(21) | Site C | 12/10/2020 | 237 days |
| ICC1(22) |  |  | 564 days |
| ADAC1 | Acer | 10/5/2021 | 261 days |
| T16 | Eastern Dry Rock | 2/11/2022 | 129 days |

### Three-dimension Terrain Model

Orthomosaics and digital terrain models (DTMs) were constructed using Agisoft Metashape Professional V1.2.6 software. Orthomosaics and DTMs followed similar protocol to other researchers and colleagues (31,32), with slight modifications. Permanent control points, galvanized nails hammered into the bedrock and epoxied at the base to the seafloor, were georeferenced using a GARMIN system on-board the vessel for image reconstruction. *A. cervicornis* cover was quantified from orthomosaics (Fig 2).

**Fig 2.**
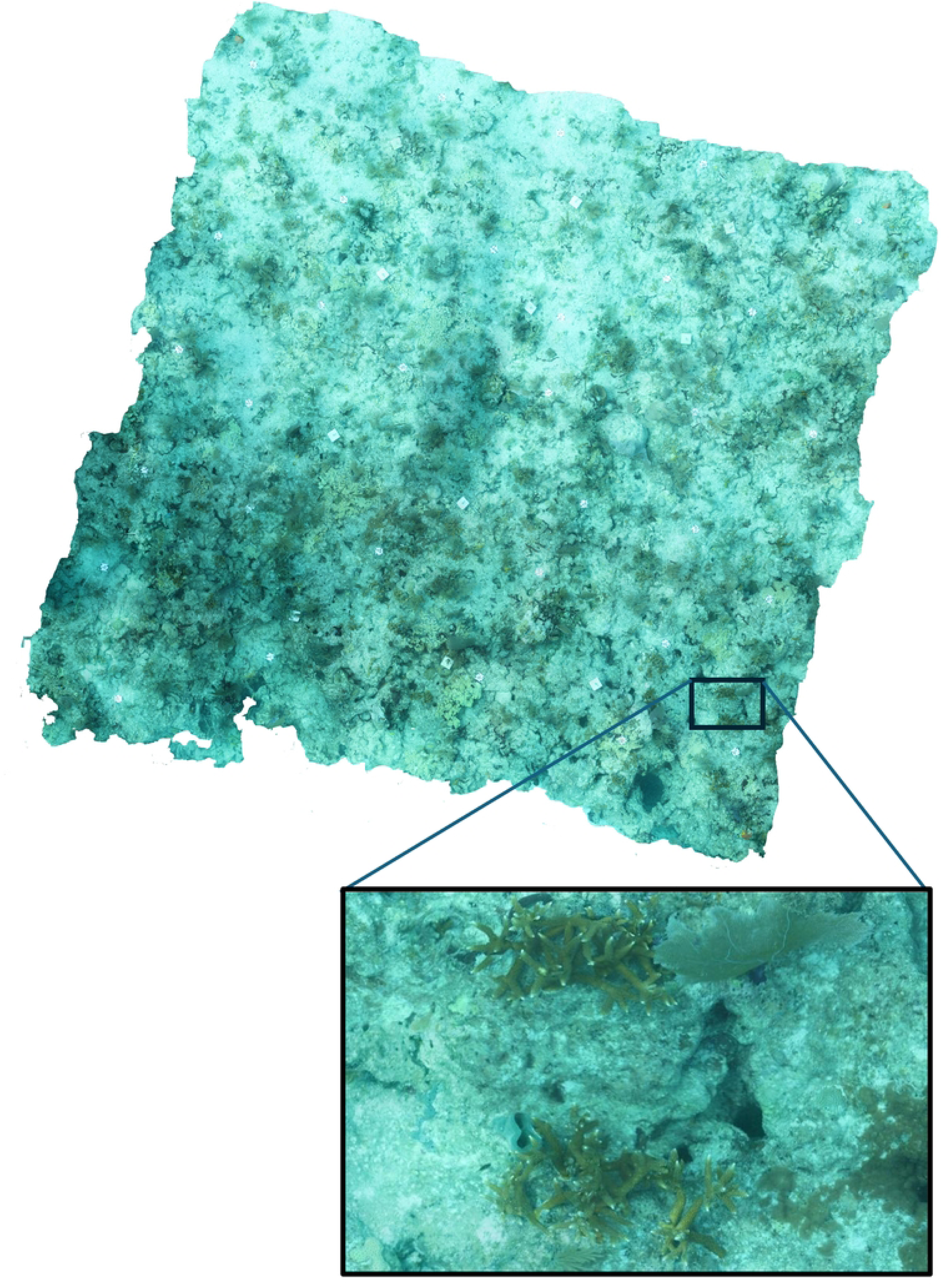
A 10 x 10m orthomosaic and a magnified section showing outplanted staghorn coral in site ICC1.

### Coral Cover

Feature class polygons of healthy coral cover and total coral cover were created. Healthy tissue cover was identified as orange/beige tissue, without any unhealthy cover (coral tissue affected by disease or covered with bacterial mats or macroalgae). Total coral cover included all skeletal tissue cover, both healthy and unhealthy. Percent healthy cover was quantified as the percentage of healthy coral cover from total coral cover. Coral cover was then classified by fragment, wild (non-outplanted), or outplanted corals. Total and healthy growth was quantified as the change in total skeletal cover and healthy tissue cover since outplantation, divided by time since outplantation. Only one site, ADAC1, had wild coral populations within the survey plot and were easily identifiable, as they were geographically distant from the outplants and much larger than the recently outplanted 3mm coral plugs. ADAC1 outplants were all singular plugs; other sites were composed of five fragment aggregates; older sites (S4, S6, T3) composed of five fragment aggregates fused together to form a mass of thickets. Average coral fragment cover at the time of outplantation was utilized to account for variations between initial size in calculating growth.

### Covariates

#### Bathymetric and Environmental Parameters

DTMs were imported into ArcGIS Pro and projected in UTM 17 zone projection with WGS84 datum. DTM resolutions were standardized (utilizing the resample tool in ArcGIS Pro) to 8mm and were georeferenced to align with the orthomosaics to ensure accurate sampling of terrain attributes around corals. To ascribe seafloor terrain attributes to coral cover, a 0.1m buffer around each coral polygon was created. Associating seafloor terrain and environmental attributes - calculated using the Spatial Analyst toolbox and Analysis toolbox in ArcGIS Pro - with total coral skeletal cover were extracted using ArcGIS Pro ModelBuilder for each coral outplant and included as covariates in the HSM (Table 2; Fig S1).

**Table 2.** Summary of spatial statistics utilized to characterize terrain and environmental attributes.

| Terrain and Environmental Attribute | Unit | Definition | ArcGIS tool | Aggregate function |
| --- | --- | --- | --- | --- |
| Depth | Meters | Distance between water surface and benthos. | Surface Information | Max |
| Slope | Degree | Benthic steepness. | Surface Parameter | Mean |
| Aspect<br>Northerness<br>and Easternness | Cosine(Degree)<br>Sine(Degree) | Direction the terrain is facing. | Surface Parameter | Mean |
| Mean Curvature | 1/Meters | Second derivative of slope. Measures convexity/denudation or concavity/accumulation of terrain. | Surface Parameter | Mean |
| Roughness | Unitless | Surface relief ratio measures a surface's unevenness. | Focal Statistics and Raster Calculator | N/A |
| Distance from Reef Edge | Meters | Apart from ADAC1, all coral outplant sites were along the edges of the reef, near drop-offs. The Euclidean distance of the *A. cervicornis* from the reef drop-off | Near | N/A |
| Distance from Coast | Meters | The Euclidean distance of the *A. cervicornis* from the nearest coast | Near | N/A |
| Distance from other *A. cervicornis* | Meters | The Euclidean distance of the *A. cervicornis* from the nearest *A. cervicornis* | Near | N/A |

#### High intensity events

Water temperature and wind data spanning 30 years were cumulatively collected from three NOAA buoys: Sand Key (January 1991 – September 2005), Sombrero Key (February 1998 – February 2008), and Key West (February 2005-January 2021). Marine heat waves (MHW) and marine cold waves (MCW) are defined herein as five consecutive days or more where water temperatures exceed the 90^th^ percentile (heat wave) or fall below the 10^th^ percentile (cold wave), based on a 30 year historical base-line (33). This day-specific definition of marine heat and cold waves accounts for extreme temperatures within a given season, which can disrupt seasonal biological processes (like reproduction). Consequently, cold waves occurring in summer months are reflected and vice versa. The duration and intensity of these events were defined by the number of days within a heat/cold wave event and the number of degrees exceeding or falling below the 90^th^/10^th^ percentile, respectively. To avoid collinearity with time, the annual average intensity of the event per duration of time throughout the corals lifetime was incorporated into the model.

High wind events were defined as any day(s) when the average daily wind speed exceeded 12.78 knots, equivalent to the average sustained wind of a tropical storm (34). Like MHW and MCS, high wind events were similarly transformed to avoid collinearity with time.

### Statistical and Model Analysis

Coral fragments were unable to characterize growth and were excluded from the analysis. All statistical analysis and models were run in R (RStudio Team, 2020). We used a single sample two-tailed Student’s t-test to determine if there was a significant difference between growth (both total and healthy) and the null hypothesis of zero growth. To evaluate differences in growth and percentage of healthy cover across sites and time, a one-way Analysis of Variance (ANOVA) was performed, with the null hypothesis of no differences among sites. If differences were observed a post-hoc Tukey’s HSD test (*p* < 0.05) was used to determine which sites differed from one another.

Five generalized linear mixed models (GLMM), operating within a frequentist statistical framework, were implemented using the glmmTMB package in R (35,36). Three models were applied to the entire dataset of observed coral outplants: one for percent healthy coral cover, one for total growth, and one for healthy growth. Dramatically different trends in percent healthy cover were observed for corals less than and greater than 2 years old, resulting in submodels being run based on these age groups (young and old coral models). Model selection took the form of excluding highly correlating covariates. No covariates were excluded in the models for total growth, healthy growth, and percent healthy cover due to adequate sample size, but submodels with reduced sample size excluded highly correlating variables. All covariates, except time, were standardized to z-score values, resulting in model coefficients indicating change in growth (cm^2^/yr) for every change in standard deviation of the environmental and terrain covariate. The percentage composition of a coral outplant with healthy tissue was doubly bound between 0% and 100% using a beta-distributed GLMM using with a logit link function. Due to logit link limitations, healthy cover equal to 0% and 100% were transformed to 0.001 or 0.999, to fall within values of 0<y<1, due to logit link limitations. Healthy and total cover models utilized a Gaussian distribution. For all models, sites were run as random effects. Model diagnostics were visually inspected and affirmed from the “Performance” package and the Akaike Information Criterion (AIC) (37). Model convergence was affirmed from the glmmTMB hessian statistic and residuals were checked for model assumptions.

## Results

After wild and fragmented corals were excluded, a total of 588 observed corals were included in the niche model. M32-3 had the highest number of coral outplants (105), and site T3 had the fewest (27) observed coral outplants. The deepest coral outplant locations were in T16 (M = 8.35m) and shallowest at T3 (M = 6.44m). The steepest coral outplant location was M32-2 (M = 37.74°) and the least steep was T3 (M = 23.99°). The least rough coral outplant locations were at site S6 (M = 0.471) and ICC1(22) had the highest roughness (M = 0.493). Qualitative observations of high surrounding roughness seem indicative of surrounding rocks and soft/other hard corals. The most concave coral outplant locations were S6 (M = -2.148 m^-1^), and the most convex locations at ADAC1 (M = 3.119 m^-1^). Covariates exhibited minimal collinearity, except for a notable correlation between distance from the reef edge and high wind events (R^2^ = -0.80). The models of young and old corals displayed high collinearity among high intensity events, as well as with distance from the reef edge and depth.

### Percent Healthy Cover Model

Temporal trends indicate healthy tissue declined with time (GLMM, *z* = -2.63, *p* = 0.008). Sites that had outplants less than 2 years old (T16, ADAC1, ICC1(21), ICC1(22)) had overall greater percentages of healthy coral cover. Among sites with coral outplants greater than 2 years old (M32-2, M32-3, S6, S4 and T3), the percentage of healthy tissue cover was lower with the lowest mean percentage of healthy tissue cover observed at T3, the oldest outplant site (Fig 3).

**Fig 3.**
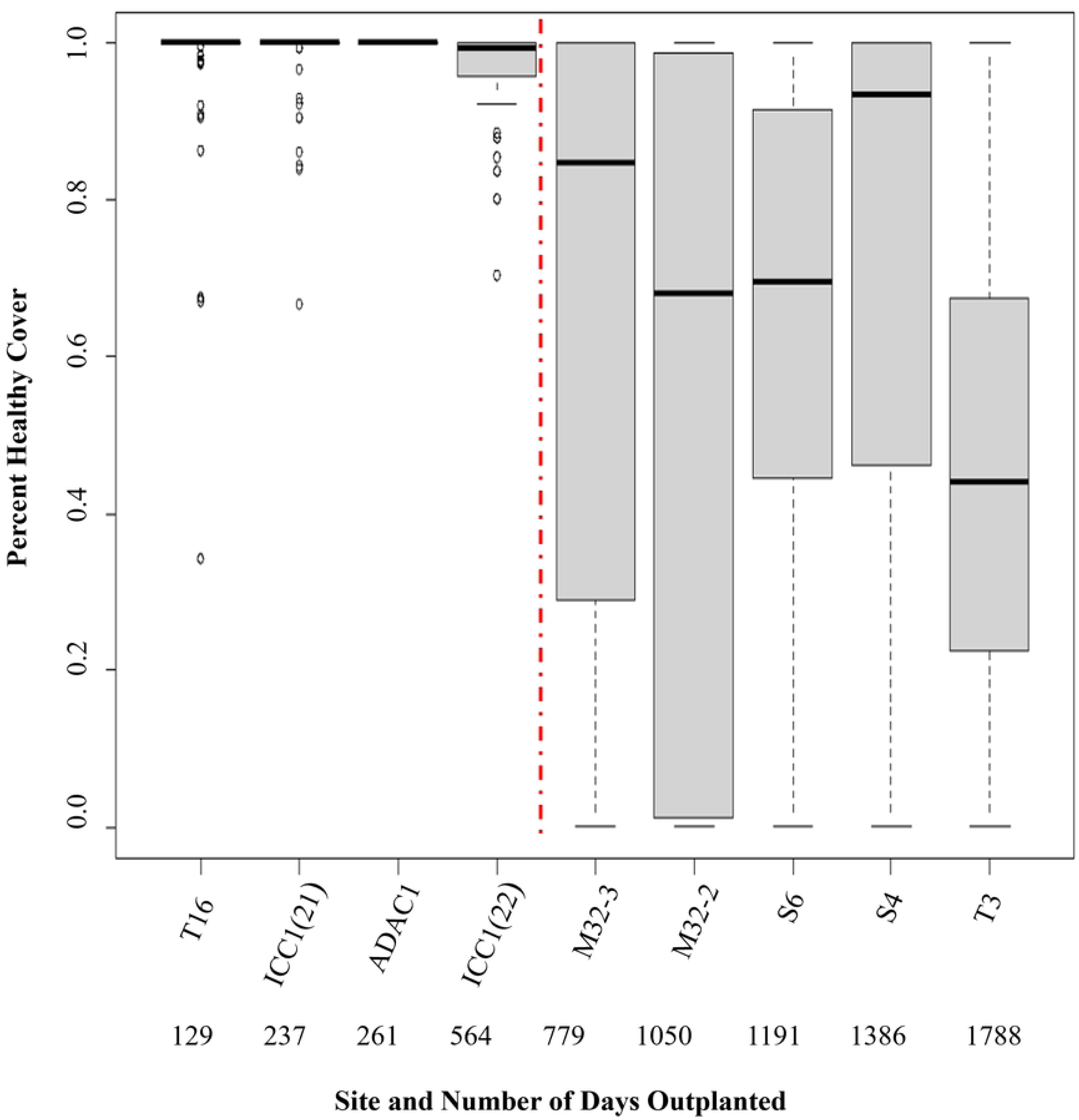
Percentage of healthy coral cover at each site. Site and associating time spent outplanted increase from left to right. Boxplots show the median, and upper and lower quartiles. Sites on the left side of the red dotted line are less than 2 years outplanted and corals on the right side are more than 2 years outplanted.

Overall, percent healthy cover was variable across and within sites. Of the 588 corals, 261 exhibited no disease or turf algae coverage and 41 exhibited no healthy cover. M32-2 had the most corals with no healthy cover. Coral outplant sites within Sand Key exhibited higher percentages of healthy cover trends than other sites with older corals (Fig 3). All coral outplant sites, apart from the most recent outplants at site ADAC1, had some corals that displayed some sign of unhealthy tissue cover. Percent of healthy cover decreased over time (GLMM, *z* = -2.63, *p* = 0.008). Sites with outplants less than 2 years old (T16, ADAC1, ICC1(21), ICC1(22)) did not differ in percent healthy coral cover and had greater percentages of healthy coral cover (ANOVA, F_8,579_= 30.89, *p* < 0.001). Post-two years of outplantation some corals observed were completely diseased and/or covered with turf algae, with no discernable healthy tissue coverage (0% healthy coral cover), and therefore assumed to be dead (Fig 3).

All models converged according to the hessian statistic and had AIC values of -1646.86 (all corals), -1118.50 (young corals), and -646.52 (old corals). Model results of young and old corals show increased distance from coast (GLMM, *z* = 2.10, *p* = 0.035), decreased distance from the reef edge (GLMM, *z* = -2.02, *p* = 0.043), increased depth (GLMM, *z* = 2.58, *p* = 0.009), and decreased roughness (GLMM, *z* = -6.28, *p* < 0.001) correlated with higher healthy coral cover (Fig 4). Submodels of young and old corals attribute the correlating relations come from older corals; no environmental variables could predict percent healthy cover in the first two years of outplantation. In contrast, increased depth (GLMM, *z* = 2.47, *p* = 0.013) and decreased roughness (GLMM, *z* = -6.20, *p* < 0.001) correlated with healthy coral cover in corals ≥ 2 yrs old (Fig 5).

**Fig 4.**
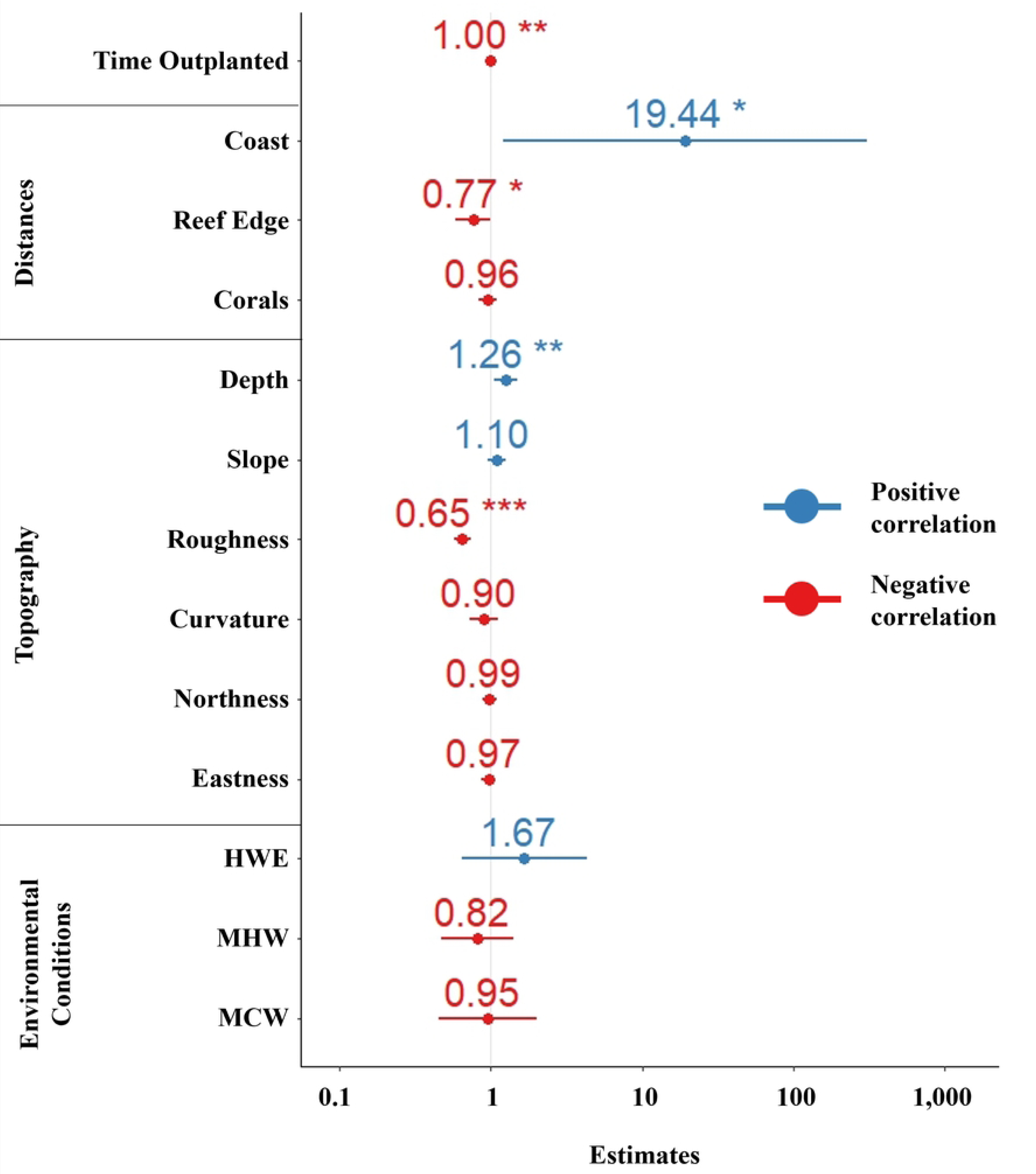
Percent coral healthy tissue model covariate estimates for each covariate for all corals. Lines depict the covariate estimates confidence intervals. Asterisks indicate the significance level of estimates.

**Fig 5.**
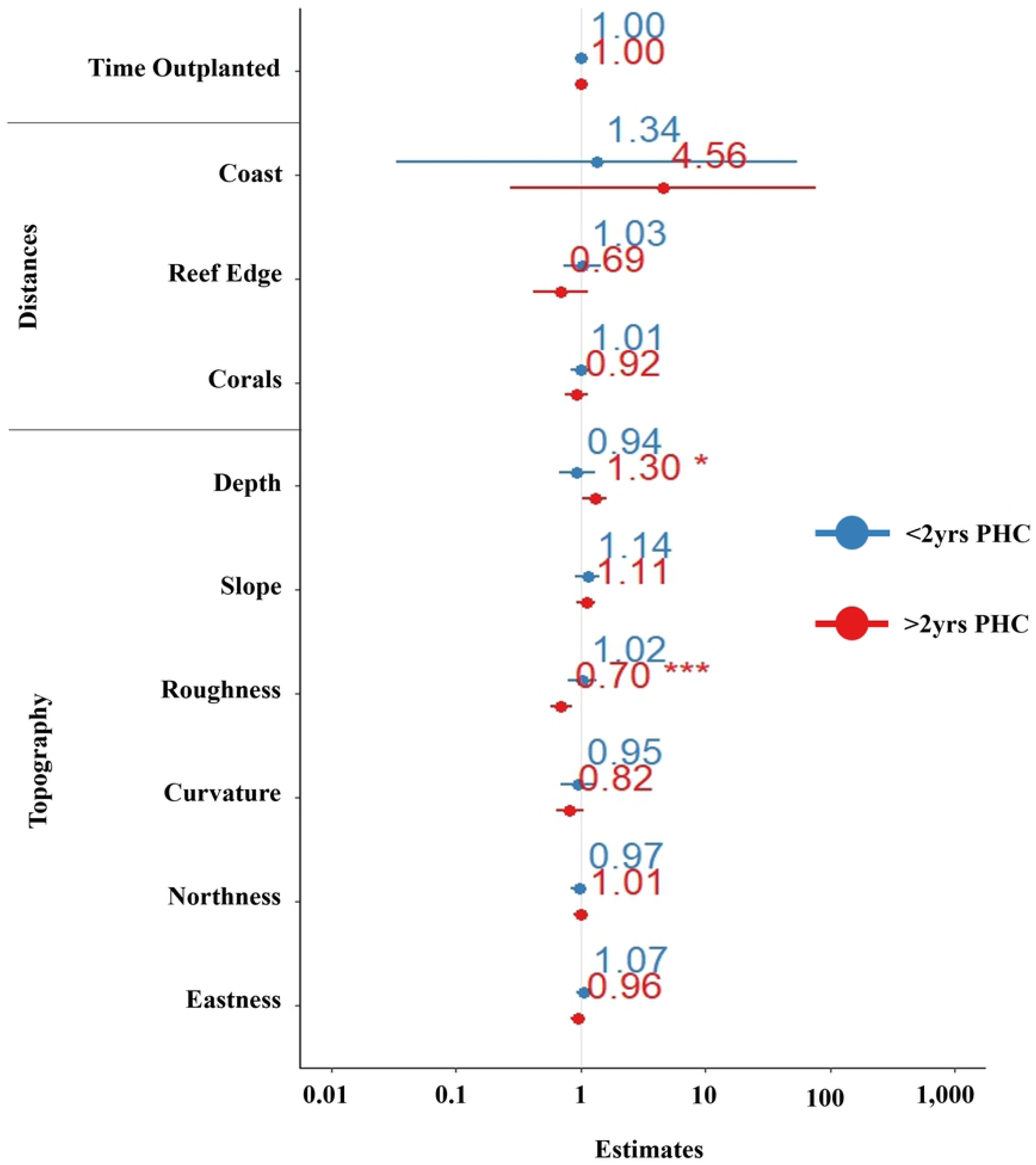
Percent coral healthy tissue model covariate estimates for each covariate for the subset models of young (<2yro) and old (>2yro) corals. Lines depict the covariate estimates confidence intervals. Asterisks indicate the significance level of estimates.

### Healthy and Total Growth Model

Outplanted corals showed total and healthy growth (t-test, t_t_ = 22.72, *p*_t_ < 0.001; t-test, t_h_ = 14.56, *p_h_* < 0.001) (Fig 6). Total and healthy growth rates varied across sites, but did not change over time (GLMM, *z_t_* = 0.97, *p_t_* = 0.33; GLMM, *z_h_* = -1.46, *p_h_* = 0.145) (Fig 6). Since the time corals were outplanted, total growth across sites averaged 26.03 cm^2^/yr (CI: 24.19-28.77 cm^2^/yr) with an average healthy growth of 16.19cm^2^/yr (CI: 14.12-18.53 cm^2^/yr).

**Fig 6.**
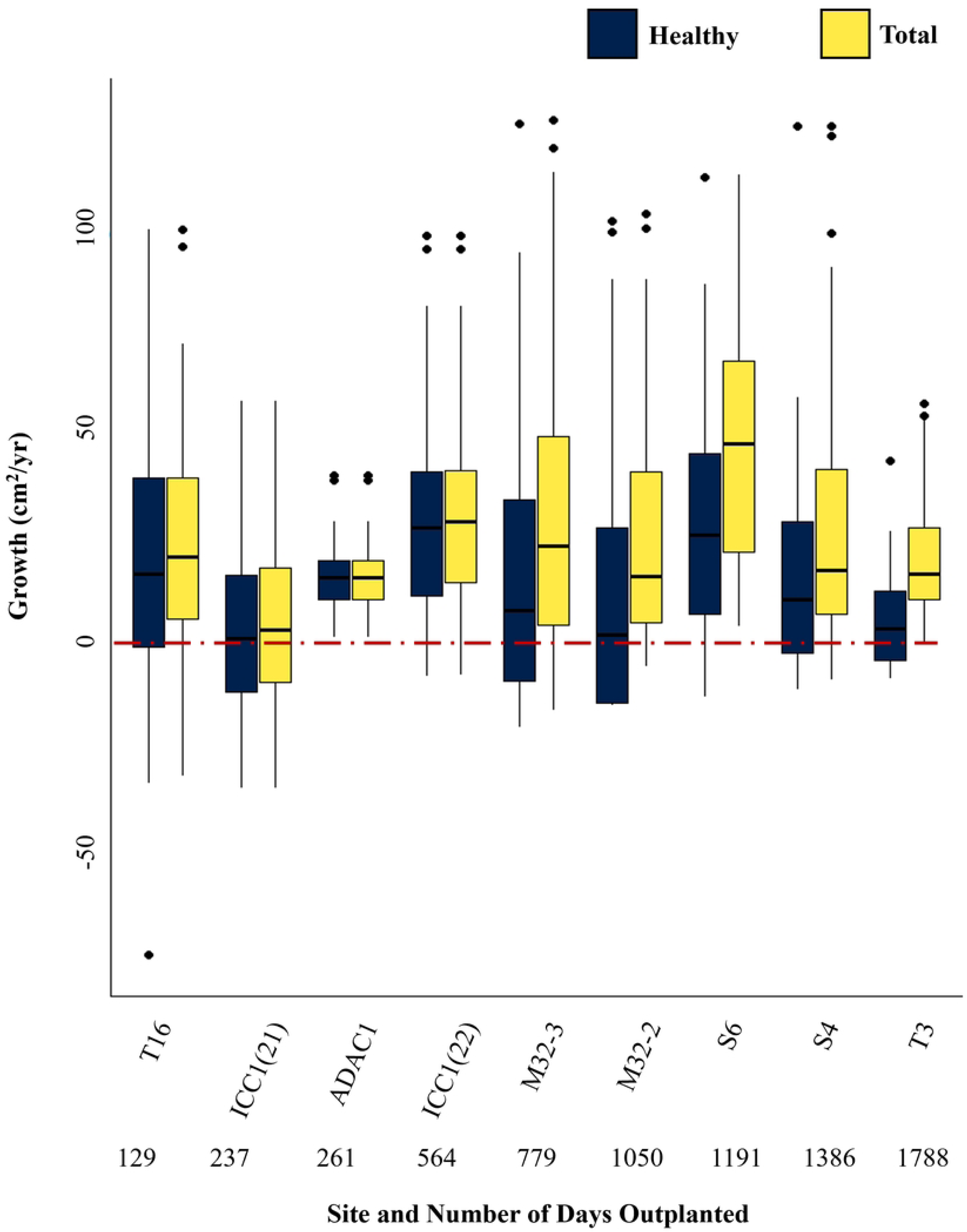
Healthy and total coral growth for all sites. The red dotted line at 0cm^2^/yr depicts where the growth rate is positive (above) or negative (below). Total and healthy growth rates could be negative due to fragmentation, reducing the coral size to smaller than its initial size. Healthy growth could be negative due to disease.

The total growth model and healthy growth model converged according to the hessian statistic and had AIC values of 4928.68 and 4955.47, respectively. Several variables correlated with higher total and healthy coral growth. This included distance from shoreline (GLMM, *z_t_* = 2.23, *p_t_* = 0.026; GLMM, *z_h_* = 2.45, *p_h_*= 0.014), increased slope (GLMM, *z_t_* = 2.27, *p_t_*= 0.023; GLMM, *z_h_* = 2.57, *p_h_* = 0.010), decreased roughness (GLMM, *z_t_* = -2.81, *p_t_* = 0.004; GLMM, *z_h_* = -4.22, *p_h_*< 0.001), decreased curvature (concavity) (GLMM, *z_t_* = -3.84, *p_t_* < 0.001; GLMM, z_h_ = -3.29, p_h_ = 0.001), increased high wind events (GLMM, *z_t_* = 2.03, *p_t_* = 0.042; GLMM, *z_h_* = 2.59, *p_h_*= 0.009), and decreased marine cold waves (GLMM, *z_t_* = -3.73, *p_t_* < 0.001; GLMM, *z_h_* = -3.16, *p_h_*= 0.0015). Increased marine heat waves correlated with lower total growth only (GLMM, *z* = -2.23, *p* = 0.025) (Fig 7).

**Fig 7.**
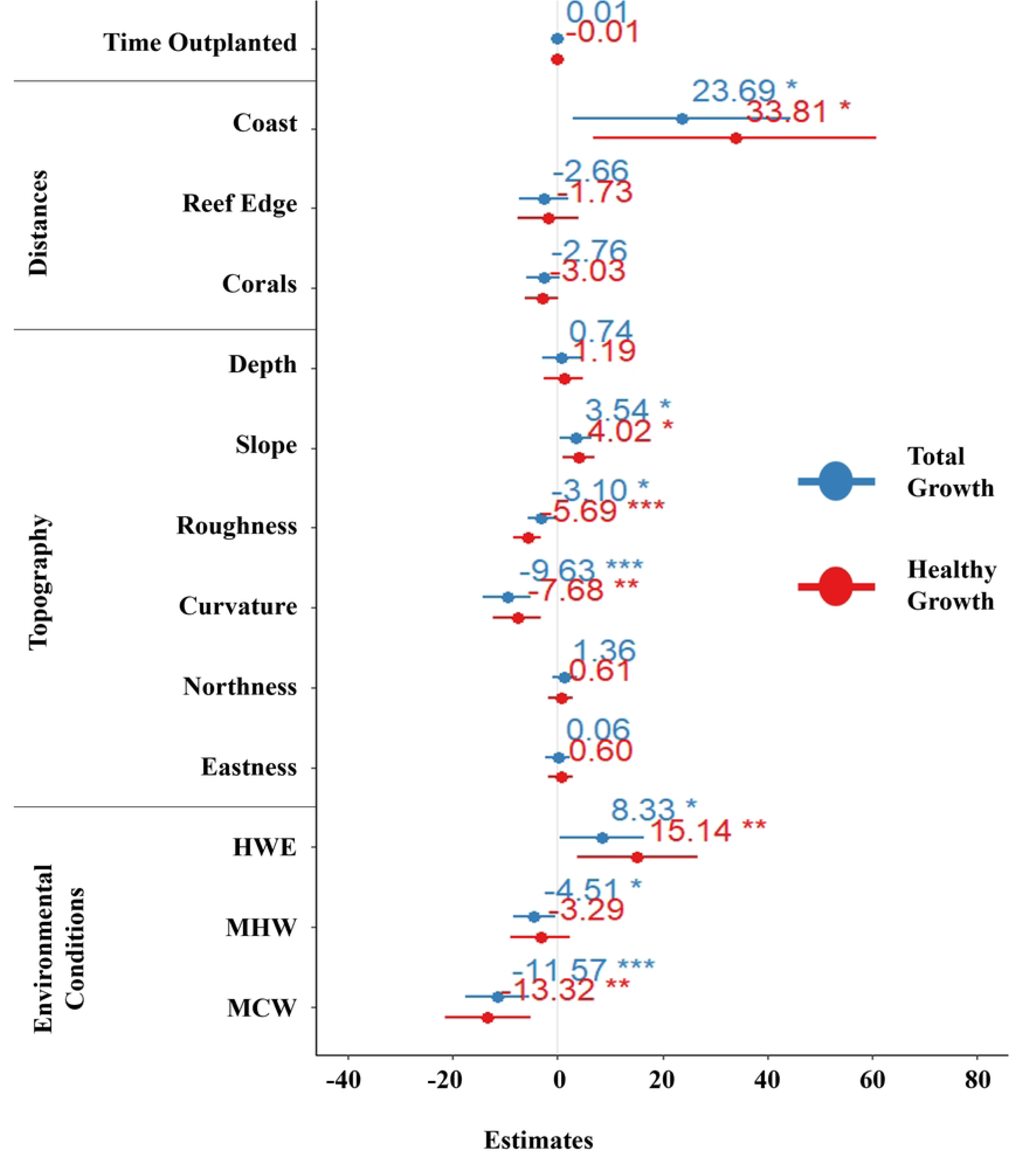
Healthy and total growth model covariate estimates for all corals. Lines depict the covariate estimates confidence intervals. Asterisks indicate the significance level of estimates.

## Discussion

Percent healthy cover, unlike growth, decreased after two years from outplantation. A possible reason for the decline in healthy coral cover after two years of outplanting may be due to the onset of spawning around 2 years of age, which likely diverts energetic resources from fighting infection to reproduction and growth (17,38,39). In contrast to results for percent healthy cover for all corals, the model for corals outplanted for less than two years was unable to identify correlating terrain attributes that positively related to percent healthy cover, emphasizing the importance of using long-term coral outplants when developing suitability models. The best predictors for high healthy coral cover over two years of age were greater depth and less roughness, indicating these environmental conditions are especially important when informing outplant locations for *A. cervicornis* reefs.

Overall, our fine-scale models suggest that depth, distance from the reef edge and the coast, and seafloor roughness correlated to percent healthy coral cover. For depth, healthy coral cover increased with depth. Scleractinian corals are sensitive to UV radiation and high temperatures, with greater temperatures and UV radiation occurring in the upper 10m (24,40,41). All our study sites were shallower than 10m. Our results revealed depth as a factor for percent cover of healthy coral, suggesting that even small differences in depth in the upper 10m of the water column may influence coral outplant success. Corals that were outplanted farther from the coastline were predicted to have greater healthy coral cover than those outplanted closer to the coastline, which supports results found by others (9). This is likely the result of the greater concentration of anthropogenic factors, such as greater thermal stress, sedimentation, and higher concentration of pollution, all of which are known to negatively impact coral health (1). Within the reefscape, corals outplanted closer to the reef edge had a greater percentage of healthy coral tissue. Reef edges are likely to have greater water flow which reduces bleaching, the build-up of sediment on the reef, and increases in mass transport (42,43), all factors that mitigate disease (12,44–46).

Interestingly and in contrast to larger scale studies and our hypothesis, healthy coral cover positively associated with decreased roughness (i.e., smoother areas). There are a number of potential reasons for these unexpected results. First, increased surrounding roughness is indicative of higher coral abundance which may facilitate contact-spread of disease (27,28,34). Second, complex environmental terrain may support a higher number of corallivores due to the greater number of available shelters. Corallivores create lesions in the coral tissues, which provides greater opportunities for disease to colonize scleractinian tissue, including *A. cervicornis* (27,47–50). Lastly, high terrain roughness and surrounding corals with rough tissue results in higher mass transfer due to an increase in micro-turbulence (27,51,52), which may lead to an increase in nutrient and gas exchange and thus a greater capacity to uptake disease-causing microbes.

Total and healthy coral growth was hypothesized to decrease over time and both growth models were assumed to perform differently. These assumptions were made due to evidence that energetic resources are diverted away from growing when approaching reproductive years or when fighting off disease (17,38,39), neither of which were evident in this study. Corals in the present study did not exhibit any decrease in growth trends, but it is important to acknowledge that this study did not include monitoring the same corals over time to establish growth trends. Further studies should monitor annual changes in coral growth or decline since outplantation. Other than MHW, covariates that significantly correlate to total growth are also significantly correlated to healthy growth in the same manner and proportion, indicating optimal conditions for growth did not change after infections.

Total and healthy growth exhibited similar correlations as percent healthy cover with distance from the coast and roughness. Additionally slope, curvature, MCW, and high wind events correlated to total and healthy growth. Less terrain roughness and increased distance from shore positively correlated with higher total and healthy growth and likely occurs for similar reasons. Steeper slopes positively correlated with healthy and total growth. Steeper slopes may indicate areas of high-water flow (53,54), suggesting that while healthy coral cover benefits from water flow, growth is dependent upon the strength of such water flow providing ample nutrients and dissolved oxygen for mass transport. Higher curvature (convex terrain) correlated with lower growth and concave terrain correlated with higher growth. Large-scale concave geomorphology has been shown to negatively correlate to coral growth, likely as a result of less water flow (55). When zooming in to observe coral biogenic morphology, concavity in coral structure causes sediment accumulation (56) and an increase ratio in polyp to surface area, causing faster food depletion (57). Intermediate of these spectrums, at fine spatial scales, convex terrain represents coral outplant locations next to encroaching soft corals and rocks, limiting water flow and competition for resources, thereby reducing mass transport (58). Convex terrain was also highly rough and therefore likely correlated negatively with growth for similar reasons.

It is not surprising that MHW and MCW negatively correlated to total and healthy coral growth. MHWs have a long-standing evidential history of adversely influencing coral success (21,40,59–62), and climate projections show MHWs are the main drivers inhibiting coral growth (63) and causing coral reef decline (2). While attention has focused on the effect of MHW over MCW, due to the expected increase in intensity and duration of MHW and decreased prevalence of MCW due to climate change (2), model results in this study revealed MCW had a greater negative impact on *A. cervicornis* total and healthy growth than MHW. One of the most severe cold-water events in the Florida Keys on record occurred in in the winter of 1976-1977 (64) and in 2010, causing mass mortalities throughout the Florida Reef Tract (8,65,66). *A. cervicornis* and other corals’ high vulnerability to cold water events raises some concerns over selective breeding and outplanting of heat tolerant corals to combat bleaching (67,68). This form of selection may leave reefs more vulnerable to the rare occurrence of cold wave events. Future selective breeding and genetic modifications should take this into account to ensure resilience under varying environmental conditions.

In contrast to other studies, our model predicts that high wind events may enhance coral growth. These events have been associated (1–3) with increased lesions (34), reduced growth (69), and increase coastal run-off (13). However, the benefits of high winds include increased wave action that can lead to higher incidences of asexual propagation through fragmentation (7,38) and increased water flow which may mitigate bleaching by bringing in cool deeper waters (70–72). Our study sites were located at depths of less than 10 m, making it likely that high wind events had a direct effect on the coral reefs. Although the impact of high wind events on corals requires further study, our findings indicate they are less of a concern than high intensity thermal and cold wave events.

### Considerations for future outplanting

Model results of curvature and roughness highlight recurring challenges with coral restoration; often coral outplanting results in unfavorable environmental conditions for optimal coral growth and survivorship. Large reef-building coral thickets can increase coral vulnerability to density induced disease spread (27,73), corallivore presence (48,74), and decreased water flow penetration into the thickets (42,75). In less rough and convex terrain *A. cervicornis* had higher growth and healthy cover trends, yet the coral itself has rough and convex morphology, resulting in the coral body building unfavorable terrain. Possible courses of actions include outplanting corals as soon as they are sexually reproductive so outplants may immediately start recruiting to the coral reef and to increase the adaptive potential of outplants to be more resilient to disease and optimize mass transport in the long-term.

Coral outplanting and monitoring capabilities can often stand at odds with maximizing restoration success. Model results within this study and others have demonstrated that the predicted suitable habitats are in deeper, cooler, waters (at least below 15m deep) that act as refuge from MHW (24,64,76), and are located farther away from the coast and its associated anthropogenic impacts (1). However, these locations are especially hard for restoration groups to regularly access and monitor coral success. Consequently, greater effort could be made to work collaboratively with restoration groups to develop habitat suitability models which identify suitable sites useful to coastal managers and practitioners.

The fine-scale digital terrain models developed from SfM can be used to inform several factors that may be useful in coral restoration areas. These include, but are not limited to, water flow (which is often used as a surrogate for the relative amount of food and oxygen delivered to species), presence of corallivores, disease, and areas that may be subject to upwelling and therefore mitigate temperature (47,77,78); all factors that help to define ecosystem health and distribution/abundance of biota (79). Model results of roughness and curvature within this study support this, where areas of high terrain roughness and convexity are primarily attributed to encroaching soft and other hard corals and rocks. The insights gained from SfM shows how higher resolution DTMs are able to bridge the gap from broader environmental conditions to more localized factors that influence organismal success (42), factors that are otherwise unavailable from satellite and sonar derived bathymetry (28,29,80,81).

Due to the insulating capacity that deeper and distant waters provide corals from climate change and anthropogenic influence, auxiliary efforts towards technological developments for easier monitoring are necessary for long-term restoration work. Current reef restoration has focused on the shallow reef crest, reef crest, spur and groove terrain, forereef terrace, and deep reef (82). These sites, apart from the deep reef, range in depth from 0.7m to 10m deep and constitute over 56,000m^2^ of restorable area (82). These areas, while accessible, are the most vulnerable to climate change, necessitating outplanting deeper and farther from the coast. Opportunities to survey coral reef growth and health more efficiently using a variety of remote sensing techniques should be explored as it can increase monitoring capabilities. Potential applications include multibeam sonar, ICESat-2 satellite monitoring, and unmanned aerial surveillance. Complementing these survey methods with HSMs and deep learning coral detection methods (83) can provide more accessible monitoring of corals in environments which may be difficult to monitor recurringly *in-situ*.

### Summary and Conclusion

As climate change progresses and environmental conditions continue to change, the available habitat for endangered species will continue to shrink. Conservation efforts to restore sensitive species populations requires attentive and rigorous research for what supports organismal success. Within modeling research, this necessitates finer-scale modeling, at the scale of meters to centimeters. While large-scale modeling is helpful towards understanding suitable habitat shifts under climate change, site-specific models are necessary to inform active restoration strategies. Structure-from-motion photogrammetry is unique from other survey methods in capturing continuous spatial data at the scale of millimeters. While SfM photogrammetry data collection is more time intensive compared to MBES, satellite, or UAS imagery data collection, the resulting data resolution is imperative to informing environmental influences on vulnerable species.

Cumulatively, our results indicate that the variance of success within the fine-scale can be accounted for and incorporated into outplant decisions. *A. cervicornis* habitat suitability models of coral growth and healthy cover inform recommendations to outplant

1. on steeper parts of the reefscape, along the reef edge
2. within marine sanctuaries, preferably in as distant locations as possible from the coast
3. in deeper waters, preferably deeper than 10m
4. where there is minimal surrounding corals and rocks

These outplant recommendations are developed with restoration group capabilities in mind, for outplant sites they are already frequenting. The creation of these models for *A. cervicornis* supports the applicability of fine-scale modeling of other endangered and vulnerable species, to support conservation efforts.

## Acknowledgement

Thank you to the managers and staff at Mote Marine Laboratory for their support with field work and commitment to restoring Florida’s reefscape. This study was supported by NOAA grant # NA20NOS4000196.

